# Saffron (*Crocus sativus*) is an autotriploid that evolved in Attica (Greece) from wild *Crocus cartwrightianus*

**DOI:** 10.1101/537688

**Authors:** Zahra Nemati, Dörte Harpke, Almila Gemicioglu, Helmut Kerndorff, Frank R. Blattner

## Abstract

*Crocus sativus* is the source of saffron, which is made from dried stigmas of the plant. It is a male-sterile triploid that ever since its origin has been propagated vegetatively. The mode of evolution and area of origin of saffron are matters of long-lasting debates. Here we analyzed chloroplast genomes, genotyping-by-sequencing (GBS) data, nuclear single-copy genes, and genome sizes to solve these controversial issues. We could place 99.3% of saffron GBS alleles in *Crocus cartwrightianus*, a species occurring in southern mainland Greece and on Aegean islands, identifying it as the sole progenitor of saffron. Phylogenetic and population assignment analyses together with chloroplast polymorphisms indicated the wild *C. cartwrightianus* population south of Athens as most similar to *C. sativus*. We conclude that the crop is an autotriploid that evolved in Attica by combining two different genotypes of *C. cartwrightianus*. Vegetative propagation prevented afterwards segregation of the favorable traits of saffron.

## Introduction

Saffron is the most expensive spice worldwide and is used since ancient times for its aroma and the ability to give dishes and textiles a golden-yellow hue (Negbi, 1999). During the last decades also medicinal properties of the plant became of interest (Abdullaev and Frenkel, 1999). The use of saffron was already documented in 3600-year-old Minoan frescos from the southern Aegean islands Crete and Santorini. At that time the plants were probably not *C. sativus* but belonged to wild saffron, *C. cartwrightianus* (Negbi and Negbi, 2002). While this latter species is an obligate outbreeding diploid, cultivated saffron is a male-sterile triploid. Both species are morphologically similar, but the stigma of the saffron crocus is thought to be longer and of darker color than that of wild saffron and the aroma is more pronounced in the cultivar. The overall similarity between both taxa was sometimes the reason to assume that *C. cartwrightianus* is the progenitor of the saffron crocus (Matthew, 1999; Negbi and Negbi, 2002). However, there are other similar species and molecular data were inconsistent regarding the contribution of possible parental taxa. Although *C. cartwrightianus* has been postulated several times as parent or one of the parental species, also *C. almehensis, C. hadriaticus, C. haussknechtii, C. mathewii, C. michelsonii, C. pallasii, C. thomasii* and *C. serotinus* have been proposed as possible parents (see Nemati et al., 2018). However, none of the analyses conducted to date have allowed for the safe inference of the parent(s), partly due to study designs that did not include the relevant species, and partly due to methodological shortcomings (Nemati et al., 2018). Additionally, the mode of evolution of triploid *C. sativus*, i.e. if it originated through autopolyploidization from a single progenitor (Brighton, 1977; Ghaffari, 1986) or allopolyploidization involving two parental species (Tsaftaris et al., 2011; Harpke et al., 2013) has been debated. In a recent study inferring phylogenetic relationships of the species of *Crocus* series *Crocus*, that is the taxonomic group to which *C. sativus* belongs, we found that *C. cartwrightianus* is the closest relative of the saffron crocus and hypothesized that no other species might have contributed to the formation of the triploid (Nemati et al., 2018).

Here we follow up on this hypothesis by analyzing genome-wide single-nucleotide polymorphisms (SNP) to see *(i*) if all alleles present in *C. sativus* can be detected within *C. cartwrightianus* or if a fraction of the saffron alleles might not be derived from this species. This should allow us to discern an auto-from an allopolyploid origin of saffron. *(ii*) SNP data are also used to find populations or areas where genetic similarity of the progenitor(s) towards *C. sativus* is highest. Depending on the species’ genetic structure, this should allow identifying the region where *C. sativus* originated. *(iii*) We test the results of SNP data by analyzing diversity of chloroplast genomes, which provide an independent source of population data. (*iv*) To understand why analyses of potential progenitor species of the saffron crocus resulted up to now in widely contradicting results we analyze allele diversity at five nuclear single-copy genes in a group of species closely related to *C. sativus*.

## Results and discussion

### Analyses of genome-wide SNP data

To see if alleles present in *C. sativus* can all be traced back to its putative progenitor *C. cartwrightianus* and to infer the geographic location of the origin of saffron, we analyzed genome-wide SNP data obtained via genotyping-by-sequencing (GBS; Elshire et al., 2011). We first processed 22 saffron individuals within the IPYRAD analysis pipeline (Eaton, 2014) to detect the loci common for 85% of the individuals, i.e. assembling a GBS reference of saffron that omits the majority of somatic present/absent mutations that might have accumulated through time within this clonal lineage as well as loci with too low or high coverage or indications that loci are not occurring in single-copy state. This reference, consisting of 6512 GBS loci (of which 1768 were heterozygous in saffron), was then used to evaluate how many of the saffron GBS alleles occur in *C. cartwrightianus* and *C. oreocreticus*. Here we consistently used 7–10 individuals out of ten *C. cartwrightianus* populations from the species’ entire distribution area (Figure 1A) and four individuals each out of two *C. oreocreticus* populations. We could place up to 90.47% of the GBS alleles occurring in saffron to the *C. cartwrightianus* populations analyzed, while 76.83% occur in *C. oreocreticus.* For the set of all 96 *C. cartwrightianus* individuals the proportion of saffron alleles was 98.58% (table 1).

**Table 1.**
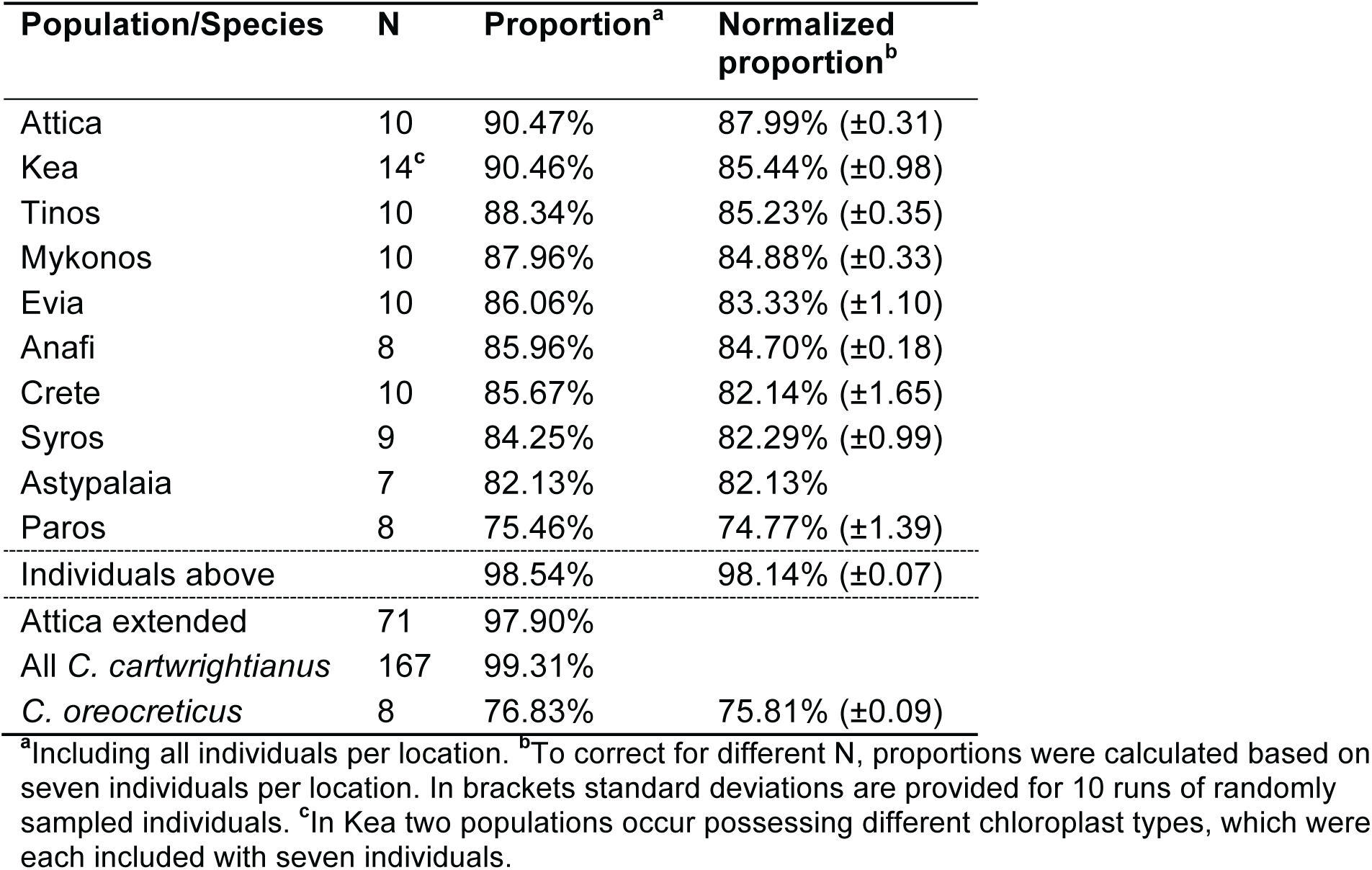
Proportion of *C. sativus* GBS alleles occurring within *C. cartwrightianus* and *C. oreocreticus*.

| Population/Species | N | Proportion<sup>a</sup> | Normalized proportion<sup>b</sup> |
| --- | --- | --- | --- |
| Attica | 10 | 90.47% | 87.99% ($\pm 0.31$) |
| Kea | 14<sup>c</sup> | 90.46% | 85.44% ($\pm 0.98$) |
| Tinos | 10 | 88.34% | 85.23% ($\pm 0.35$) |
| Mykonos | 10 | 87.96% | 84.88% ($\pm 0.33$) |
| Evia | 10 | 86.06% | 83.33% ($\pm 1.10$) |
| Anafi | 8 | 85.96% | 84.70% ($\pm 0.18$) |
| Crete | 10 | 85.67% | 82.14% ($\pm 1.65$) |
| Syros | 9 | 84.25% | 82.29% ($\pm 0.99$) |
| Astypalaia | 7 | 82.13% | 82.13% |
| Paros | 8 | 75.46% | 74.77% ($\pm 1.39$) |
| Individuals above | | 98.54% | 98.14% ($\pm 0.07$) |
| Attica extended | 71 | 97.90% |  |
| All *C. cartwrightianus* | 167 | 99.31% |  |
| *C. oreocreticus* | 8 | 76.83% | 75.81% ($\pm 0.09$) |

<sup>a</sup>Including all individuals per location. <sup>b</sup>To correct for different N, proportions were calculated based on seven individuals per location. In brackets standard deviations are provided for 10 runs of randomly sampled individuals. <sup>c</sup>In Kea two populations occur possessing different chloroplast types, which were each included with seven individuals.

**Figure 1.**
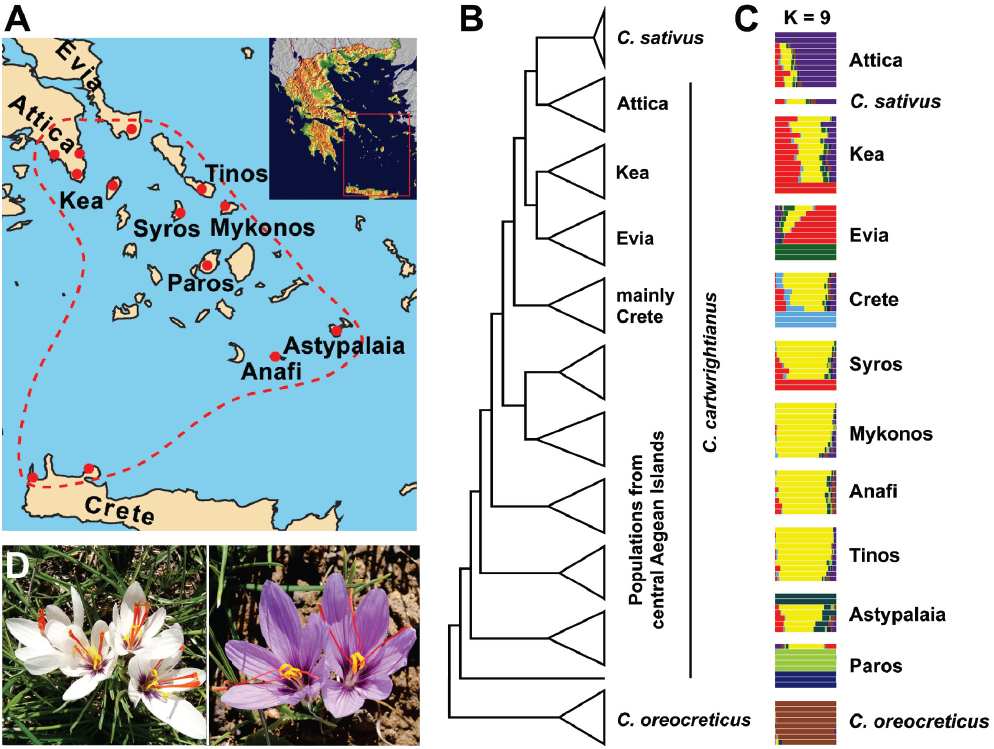
Distribution and phylogenetic relationships of *C. cartwrightianus* with regard to *C. sativus.* (A) Map providing the distribution area of *C. cartwrightianus* (dashed line). Red dots indicate the collection sites of populations included in the study. (B) Scheme summarizing the topology of two most parsimonious trees based on genome-wide DNA data of a genotyping-by-sequencing analysis of 114 individuals. *Crocus oreocreticus* was defined as outgroup. For clades within *C. cartwrightianus* the geographical origins of the samples are given. (C) Result of a Bayesian population assignment analysis for K = 9. (D) Individuals of *C. cartwrightianus* (left) and *C. sativus* (right). The following figure supplements are available for figure 1: **Figure supplement 1 – 3** **Figure supplement 6**

As *C. sativus* individuals are genetically very similar (Busconi et al., 2015) and this is also the case for the individuals we studied, we included ten of them in phylogenetic analyses based on the GBS dataset. In all analyses the samples of *C. sativus* grouped within *C. cartwrightianus* populations from the northern part of the species’ distribution range and formed the sister group of the Attic individuals (Figure 1B, Figure S1 and S2; figures and tables indicated by “S” are available as Supplementary Files online). GBS-based genetic distances in *C. cartwrightianus* are an order of magnitude higher than in saffron (up to 0.41% vs. 0.032%), and also within the *C. cartwrightianus* populations genetic diversity is comparatively high (e.g., for Attica 0.29%), This is also reflected in the respective branch lengths of the phylogenetic trees (Figure S1 and S2). Differences in amount of genetic diversity between both species were also reported by Larsen et al. (2015). For a Bayesian population assignment analysis (Pearse and Crandall, 2004) we identified an optimum of nine groups of genotypes (K = 9) including *C. oreocreticus*. Here saffron is assigned to the individuals from Attica (Figure 1C, Figure S3). However, we did not find a single *C. cartwrightianus* individual that is identical, or even most similar, to saffron but could only identify the entire Attica population as closest relatives of the cultivar, possessing 90.47% of the saffron GBS alleles. This correlation also holds after correction for sample size (Table 1).

To see if all alleles occurring in *C. sativus* can be found in Attic *C. cartwrightianus* individuals we increased our sample from Attica to 71 individuals, which we genotyped by GBS. We could place 97.60% of the saffron GBS alleles in this set of Attic samples and together with the individuals from the other areas they accounted for 99.31% of the saffron alleles (Table 1). Thus, only 0.69% of them could not be assigned to any of the included *C. cartwrightianus* individuals. This clearly indicates that only wild saffron contributed genetic material to saffron. Otherwise a much higher proportion of saffron GBS alleles should occur that are different from the ones found in *C. cartwrightianus*.

### Analysis of chloroplast genome diversity

BLAST searches revealed that none of the 6512 GBS loci of saffron we used were derived from the chloroplast or mitochondrial genome. Therefore, we analyzed DNA differences in the maternally inherited chloroplast to base our conclusions also on a marker type that is independent from the nuclear GBS data. To obtain initial information about potentially informative loci we used genome skimming (Straub et al., 2012), which is based on low-coverage whole-genome shotgun (WGS) sequencing, and assembled the chloroplast genomes (Figure S4) of two *C. cartwrightianus* individuals from the southern (Crete) and northern (Attica) borders of the species’ distribution area. The alignment of both chloroplast genomes had a length of 150,942 nucleotides and showed 99.97% identity. Among the differences was an 84 base pair (bp) deletion in the *trn*S–*trn*G intergenic spacer in the Attic individual that we found also in an individual of *C. sativus*. We used this marker as sequence characterized amplified region (SCAR) to screen for chloroplast differences in *C. sativus* and *C. cartwrightianus* individuals from all populations. We found that all *C. sativus* individuals possess the short allele while in *C. cartwrightianus* it occurs only in Attica and in one out of two populations from the island of Kea (Figure 2). Kea is directly adjacent of Attica (Figure 1A) and was connected to the mainland repeatedly when Quaternary sea levels dropped (Lambeck, 1996). This may have allowed gene flow between the crocus stands of these areas. In all other populations of *C. cartwrightianus* only the longer allele was detected (Figure 2).

**Figure 2.**
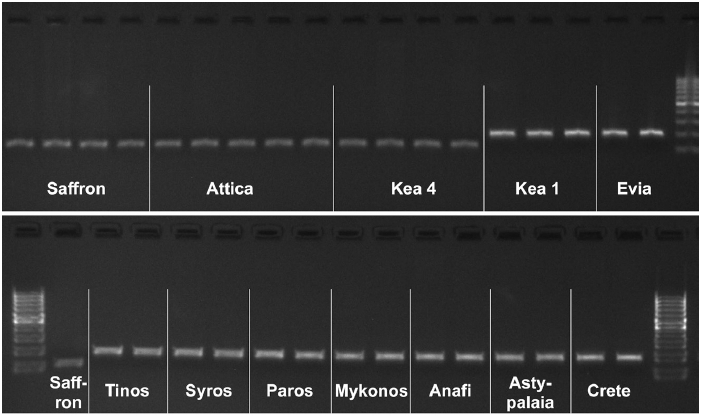
Gel photos of exemplary samples of PCR amplicons for the distribution of an 84-bp deletion in the chloroplast *trn*S-*trn*G intergenic spacer of *C. cartwrightianus* and *C. sativus*. The shorter allele (119 bp) occurs only in Attica, one population from Kea and in *C. sativus*, while all other populations of *C. cartwrightianus* possess the longer allele (203 bp). The following figure supplement is available for figure 2: **Figure supplement 4**

To further compare the chloroplast sequences of saffron and Attic *C. cartwrightianus*, the chloroplast genome of a *C. sativus* individual was assembled. Comparing both sequences we found them to be identical except for an additional adenine occurring at position 1509 of the gene for the beta subunit of the acetyl-CoA carboxylase (*acc*D) in *C. sativus*. This mutation results in a stop codon terminating the *acc*D coding region three amino acids earlier than in the wild type. We re-sequenced the relevant region of the *acc*D gene in 60 individuals of Attic *C. cartwrightianus* but could not detect the mutated variant in these samples. Still, the sole occurrence of the deletion in the *trn*S–*trn*G region in the chloroplast genome in Attic and Kean *C. cartwrightianus* individuals clearly places the maternal parent of the saffron crocus in this area of Greece and, thus, is in accord with the GBS data.

### Analysis of genome size in *C. cartwrightianus*

To be able to infer the mode of origin of the triploid, i.e. if it evolved through a cross between a di- and a tetraploid parent or through the combination of a reduced with an unreduced gamete within diploids, we collected leaves from 100 *C. cartwrightianus* individuals in the Attica area, dried them in the field in silica gel, and analyzed genome sizes for these individuals in the lab by flow cytometry. We were able to obtain results for 91 individuals. We found that all of them have a 2C genome size of 7.06 ±0.09 pg (SD). This value was also observed in individuals with chromosome counts of 2*n* = 2*x* = 16. From this we conclude that *C. cartwrightianus* in Attica is diploid and that tetraploid plants are not frequent. This is in accord with the observation that *C. sativus* is a clonal lineage that originated most probably only once, indicating that it was a rare event. If tetraploid individuals would occur regularly within the diploid *C. cartwrightianus* populations, triploid plants should arise over and over again through crosses between both ploidy levels. Over time such continuous input of triploids into the saffron gene pool should have broadened the genetic diversity occurring in saffron. This is, however, not the case as saffron is genetically rather uniform (Nemati et al., 2014; Busconi et al., 2015).

### Analysis of nuclear single-copy loci

Taking into account the high genetic diversity in *C. cartwrightianus*, we hypothesized that this could have influenced the outcome of earlier phylogenetic studies where *C. cartwrightianus* and *C. sativus* were included but did not result as sister species. We used DNA sequences of five single-copy genes that were amplified from *C. cartwrightianus, C. sativus* and their four closest relatives *C. hadriaticus, C. oreocreticus, C. pallasii* and *C. thomasii* (Nemati et al., 2018), which all share the same chromosome base number of *x* = 8. Where initial direct sequencing provided evidence for the presence of more than one copy of a gene within an individual, amplicons were cloned and six clones per individual were sequenced. Phylogenetic analyses of the DNA sequences of the five genes (Figure S5) revealed in all cases that alleles occurring in different species were not completely sorted according to their species affiliation. This phenomenon, referred to as incomplete lineage sorting (Maddison, 1997), is often found among closely related species (Jakob and Blattner, 2006; Brassac and Blattner, 2015). Even the different alleles or homeologs detected within the *C. sativus* individual were rather diverse and could group in different clades in the gene trees. Thus, the high genetic diversity in *C. cartwrightianus* is not restricted to non-coding parts of the genome (Larsen et al., 2015) but concerns also the gene space of this species, which could have rather diverse allelic constitutions.

## Conclusions

Earlier molecular studies of saffron evolution did not arrive at clear results regarding the parental species of *C. sativus* or the area of origin of saffron. The main reason seems that they did not take into account the high intra-specific genetic diversity present in *C. cartwrightianus*. Depending on the individual(s) studied and the marker region used, the resulting phylogenetic trees might reflect nearly arbitrary relationships (Figure S5). In contrast, our GBS data were based on an exhaustive collection of *C. cartwrightianus* populations and clearly place the *C. sativus* individuals as sister of Attic *C. cartwrightianus*. Possible reasons for the sister group position instead of grouping within the Attic population is most probably the triploid and clonal nature of *C. sativus* that, as a group, has therefore a unique character combination that is in this way not present in any individual of *C. cartwrightianus*. Still, overall frequencies of GBS alleles (Table 1) and also chloroplast data (Figure 2) support the saffron origin within the Attica/Kea region with closest similarity of the saffron crocus to the *C. cartwrightianus* plants occurring in Attica (Figure 1B, 1C).

When collecting leaves of Attic *C. cartwrightianus*, we evaluated the populations for the presence of the important traits typical for the saffron crocus. We recognized the bunchy growing habit, very long stigmas, stigmas of rather dark red color (Figure 1D, Figure S6), and also stigmas with the specific taste and aroma of saffron, particularly in the southern part of this area. However, we did not find plants combining all these traits within single individuals in the same way as saffron. As genetic diversity is high in *C. cartwrightianus* and the species is an obligate outbreeder, it is unlikely to find in today’s individuals regularly the exact allele combination characteristic for triploid *C. sativus*, as allele composition is constantly jumbled by genetic recombination. This is also apparent in the diverse karyotpyes of *C. cartwrightianus*, where saffron-specific chromosomes can be found, although they are not united within single *C. cartwrightianus* individuals (Schmidt et al., 2019).

From Minoan frescos it is clear that more than 3600 years ago humans already used wild saffron in the southern Aegean. The first clear indication for the cultivation of triploid saffron can be found in *Historia Plantarum* (350 BC–287 BC) where Theophrastus described the plant as being propagated by corms (Negbi and Negbi, 2002). Our GBS data point to the small Greek region south of Athens as the place where saffron evolved. We assume that sometime in between 1600 BC and 350 BC a triploid *C. cartwrightianus* cytotype originated in Attica and was selected by humans. They must have realized that they have a highly aromatic and stable type at hands that keeps the valuable properties of saffron through time and (vegetative) generations. A bit surprising is the fact that the main growing regions for saffron are today found clearly outside the distribution area of *C. cartwrightianus*, i.e. in the western Mediterranean (Spain, Morocco) and western Asia (Iran to northern India). While *C. cartwrightianus* is restricted to the Mediterranean vegetation zone, saffron cultivation happens mostly in much drier regions and at higher elevation. Thus, we assume differences in climate requirements between both species. This could indicate an ecologic niche shift due to polyploidization. We cannot yet determine how much triploidy influences the development of the typical traits of saffron or if the right allele combination in a diploid might provide similar characteristics. Still, the clarification of the mode of evolution of *C. sativus* now opens up a route for overcoming the low genetic diversity present in the saffron crocus, as it will foster new saffron genotypes to be created from different *C. cartwrightianus* individuals.

## Materials and methods

### Taxon sampling

We included 197 individuals in the genotyping-by-sequencing (GBS) study, 22 belonging to *C. sativus*, 167 to *C. cartwrightianus* and eight to *C. oreocreticus* (Table S1). The sampling of *C. cartwrightianus* covered its entire distribution range (Figure 1A). *Crocus oreocreticus*, the closest relative of *C. cartwrightianus* and *C. sativus* (Nemati et al., 2018), is endemic to Crete, where it was collected in two different populations with four individuals each. In addition we used single individuals of the other *Crocus* series *Crocus* species sharing the chromosome number of 2*n* = 2*x* = 16 with *C. cartwrightianus*, which were often named as parents of triploid *C. sativus* (2*n* = 3*x* = 24). These are *C. hadriaticus, C. pallasii* and *C. thomasii*, which were included in screens of allelic diversity at nuclear single-copy genes. Voucher information for the analyzed taxa is provided in Table S1.

### DNA extraction and PCR reactions

Extraction of genomic DNA was carried out using DNeasy Plant DNA Extraction Kit (Qiagen) from about 10 mg of silica-dried leaf material according to the protocol of the manufacturer. DNA concentration and quality were afterwards checked on 0.8% agarose gels.

To obtain nuclear single-copy marker regions with high variability in *Crocus*, we used contigs derived from the assembly of low-coverage next-generation sequencing (Illumina HiSeq platform; see below) of *Crocus cartwrightianus*. Potential nuclear single-copy genes and their intron-exon borders were identified using the PLAZA v2.5 and v3 platform (Van Bel et al., 2012; Vandepoele, 2017).

We selected five nuclear single-copy genes (Table S2), which all were heterozygous in *C. sativus*, and PCR amplified them in five *Crocus* species closely related to saffron. PCR was performed with 1 U Phusion High-Fidelity DNA Polymerase (Thermo Scientific) in the supplied Phusion GC Buffer, 200 μM of each dNTP, 0.5 μM of each primer, and about 20 ng of total DNA in 50 μl reaction volume in a GeneAmp PCR System 9700 (Perkin-Elmer). Amplification was performed with 3 min initial denaturation at 95 °C and 35 cycles of 30 s at 95 °C, 25–60 sec at the marker-specific annealing temperature (table S2) and 30 s at 70 °C, followed by a final extension for 8 min at 70 °C. PCR products were purified on a 1% gel and extracted using QIAquick Gel Extraction Kit (Qiagen) following the manufacturer’s protocol, and eluted in 30 μl water. Both strands of the PCR products were initially directly sequenced with Applied Biosystems BigDye Terminator technology on an ABI 3730xl automatic DNA sequencer using the primers from PCR amplifications. When direct sequencing revealed polymorphic sequence positions or length differences, PCR products were cloned into the pGEM-T Easy vector (Promega) and six clones per individual were sequenced with Templi-Phi DNA Sequencing Template Ampflication Kit (Amersham Biosciences).

### Library preparation and next-generation sequencing

To obtain genome-wide single-nucleotide polymorphisms (SNPs), genotyping-by-sequencing (GBS) analyses (Elshire et al., 2011) were conducted for 197 individuals. For the library preparation 200 ng of genomic DNA were used and cut with the two restrictions enzymes *Pst*I-HF (NEB) and *Msp*I (NEB). Library preparation, individual barcoding, and single-end sequencing on the Illumina HiSeq 2000 followed Wendler et al. (2014).

For WGS sequencing of *C. cartwrightianus* from Crete and Attica 1-2 μg DNA were used. Library preparation was carried out as described by Meyer and Kircher (2010) for *C. cartwrightianus* from Crete. Library preparation for *C. cartwrightianus* from Attica was done according to the Illumina TruSeq DNA library preparation protocol following the manufacturer’s recommendations. DNA was covarized to generate fragments of on average 300-400 bp length for the Crete and 400-600 bp for Attic *C. cartwrightianus*, followed by adaptor and barcode ligation. For *C. cartwrightianus* from Crete an additional 8-kb mate-pair library was generated. The libraries were size-selected with a SYBR Gold stained electrophoresis gel. Fragment size distribution and DNA concentration were evaluated on an Agilent BioAnalyzer High Sensitivity DNA Chip and using the Qubit DNA Assay Kit in a Qubit 2.0 Flurometer (Life Technologies). Finally the DNA concentration of the libraries was checked by a quantitative PCR run. Cluster generation on Illumina cBot and paired-end sequencing (*C. cartwrightianus* from Crete: 2×100 bp; *C. cartwrightianus* from Attica: 2×250 bp) on the Illumina HiSeq 2000/2500 platform followed Illumina’s recommendation and included 1% Illumina PhiX library as internal control. The 8-kb mate-pair library of *C. cartwrightianus* from Crete was sequenced using 20% of a lane on the Illumina MiSeq platform generating 2×250 bp paired-end reads.

Barcoded reads were de-multiplexed using the CASAVA pipeline 1.8 (Illumina). The obtained raw sequencing reads were quality checked and over-represented, i.e. clonal reads were detected with FASTQC (Andrew, 2010). Adapter trimming of sequence reads was performed with CUTADAPT (Martin, 2011) and reads shorter than 60 bp after adapter removal were discarded.

### Genome assembly of whole-genome shotgun data and chloroplast genomes

*De novo* assembly of WGS sequences of *C. cartwrightianus* from Crete with 210 million quality-filtered read pairs was performed in CLC v4.3.0 (CLC bio) with a minimum length for assembled contigs of 500 bp. NCBI BLAST v2.2.28+ searches were used to check for bacterial contaminations in the sequence reads and to identify plastid-derived contigs. Scaffolding with SSPACE v3.0 (Boetzer et al., 2011) was performed with a minimum number of 100 linked reads to compute a scaffold.

Plastid scaffolds were identified by a BLAST search. The scaffolds were then mapped to the chloroplast genomes of *Beta vulgaris* (GenBank accession number KR230391), *Haloxylon persicum* (KF534479) and *Iris gatesii* (KM014691). GAPFILLER v1.10 (Boetzer and Pirovano, 2012) was used to fill gaps. Proper pairing of reads was checked by mapping the original reads against the obtained *C. cartwrightianus* plastid genome using GENEIOUS R10.2.3 (Biomatters Ltd.) and checked manually.

Illumina sequencing with lower coverage produced 123.6 million (61.8 million paired end) quality-filtered reads for *C. cartwrightianus* from Attica. For *C. sativus* 74.4 million (37.2 million paired end) quality-filtered reads from a WGS experiment were provided by Thomas Schmidt and Tony Heitkam (Molecular and Cell Biology, University of Dresden, Germany) and assembled as before.

The chloroplast genome of *C. cartwrightianus* from Crete obtained by *de novo* assembly was used as a reference sequence for chloroplast genomes of *C. cartwrightianus* from Attica and *C. sativus*. Annotation of chloroplast genomes was performed in GENEIOUS and edited manually.

### Chloroplast polymorphisms screening

A sequence part of the chloroplast *trn*S(GCU)–*trn*G(UCC) intergenic spacer, where an 84-bp deletion was observed in *C. sativus* and *C. cartwrightianus* from Attica, as well as a 1-bp insertion in the *acc*D gene in *C. sativus* at position 1509 were confirmed by Sanger re-sequencing (Table S2). Screening of the distribution of the 84-bp deletion in the *trn*S-*trn*G intergenic spacer was conducted by PCR amplification of the respective chloroplast region in the set of 96 *C. cartwrightianus* individuals covering the populations from the entire distribution area of the species (table 1) plus the *C. sativus* individuals. Size determination of the PCR products was done on 1.4% agarose gels against a 100-bp DNA ladder and always including one saffron amplicon as reference for the short allele.

### Processing of genotyping-by-sequencing data

To have a particular focus on loci present in *C. sativus* we first generated a loci reference file using IPYRAD v0.7.19 (Eaton, 2014). An assembly of the GBS data of 22 saffron individuals was done *de novo*. The minimal number of samples per locus was set to 18, the clustering threshold of reads within and between individuals was set to 0.9. The maximum ploidy level was appointed as triploid. For the other parameters the default settings of parameter files generated by IPYRAD were used. The locus file generated by IPYRAD was converted into a FASTA file with 6512 sequences corresponding to the number of obtained loci.

Additional IPYRAD analyses were run, using either 126 or 197 individuals and the reference-based assembly of IPYRAD. The clustering threshold of reads within and between individuals was set to 0.9. The maximum ploidy level was appointed as triploid. For the other parameters the default settings of parameter files generated by IPYRAD were used except for the maximal number of indels, which was increased to 15. The minimal number of samples per locus was set to 18 to generate the output file used for the determination of the proportion of SNPs *C. sativus* shares with *C. cartwrightianus* and *C. oreocreticus*. For Bayesian assignment as well as the phylogenetic analyses, the minimal number of samples per locus was set to 55. VCFTOOLS 0.1.14 (Danacek et al., 2011) were used to filter out the genotypes with a depth below six.

### Determining the proportion of shared GBS alleles

The number of shared GBS alleles between the species and populations was inferred using PopGenReport (Adamack and Gruber, 2014) in R 3.5.0. In principle, the filtered vcf file was first converted into a genind object, then concatenated with the information about the sample origin and converted into a genpop object. The genpop file directly includes the allelic counts per variable position, which we here simply refer to as SNPs. SNPs not present and SNPs potentially originating by autosomal mutation in saffron (present in less than four counts in saffron) were excluded. Counting and calculating percentages were conducted in Microsoft EXCEL v14 after transposing the genpop matrix in R. In addition to the dataset where we used all individuals per population we also calculated the proportion of shared GBS alleles for seven (lowest number of individuals among our population samples) randomly chosen individuals for the populations where higher individual numbers were available. For this normalized dataset the included individuals were permuted ten times and the percentage of shared GBS alleles was averaged over the ten runs.

### Phylogenetic and population genetic analyses

Forward and reverse sequences for the five nuclear and two chloroplast marker regions were manually checked, edited where necessary, and assembled in one sequence for each locus and individual. In cases where cloning revealed the presence of different alleles, cloned sequences were assigned to different haplotypes or consensus sequences were generated for cloned sequences differing by only one substitution.

Phylogenetic analysis using maximum parsimony (MP) was performed in PAUP* 4a161 (Swofford, 2002) using either the branch-and-bound algorithm (in cases when fewer than 15 sequences of the single-copy loci occurred in a dataset) or the heuristic search algorithm with TBR branch swapping for the concatenated GBS-derived sequences and single-copy loci with 15 and more sequences. Gaps were treated as missing data and we used 1000 random-addition-sequences (RAS) to construct the starting trees in the heuristic search for the GBS data to avoid suboptimal tree-islands. Bootstrap support values were obtained by 500 bootstrap re-samples using the same settings as before except that we excluded RAS. Maximum-likelihood analyses were conducted in RAxML v. 8.0.0 (Stamatakis, 2014) using GTRGAMMA and 100 parsimony starting trees. The number of bootstrap re-samples was set to 500.

Pairwise genetic distances within the GBS dataset were calculated in PAUP* as uncorrected (“p”) distances for the *C. sativus* individuals and the *C. cartwrightianus* individuals within and among populations. In both species genetic diversity was rather uniformly distributed among the individuals.

Bayesian assignment analysis for the GBS data was carried out using the LEA package (François, 2016) in R 3.5.0 in an initial run for K = 1 to K = 20. The lowest entropy here was observed for K = 9. Therefore, for K = 2 to K = 14 Bayesian assignment analysis was run with 20 repetitions. The run and repetition with lowest cross entropy was identified and selected. Additionally, the populations of the sampling data were read in using data.table (Dowle and Srinivasan, 2018). The *Q*-matrix obtained by LEA, including the ancestral assignment frequencies, was sorted in R with Tidyverse (Wickham, 2017) according to the population the samples belong to. Plots were arranged with the gridExtra R package (Auguie, 2017).

### Genome size determinations in Attic *C. cartwrightianus*

To infer genome sizes and ploidy level of Attic *C. cartwrightianus* individuals we collected a leaf from each of 100 individuals in the field and dried them in silica gel. Afterwards the leaves were transported to the lab and genome sizes were measured with a CyFlow Space (Sysmex Partec) flow cytometer against a *Vicia faba* size standard (26.5 pg 2C) using DAPI as staining reagent and essentially following the procedure described by Jakob et al. (2004).

## Acknowledgements

We like to thank the Greek authorities for providing permits for plant collections, I. Faustmann, C. Koch, B. Kraenzlin and P. Oswald for help with plant cultivation and lab work, T. Schmidt and T. Heitkam for access to saffron whole-genome shotgun data, A. Himmelbach and S. König for performing Illumina sequencing, and H. Poskar and J. Brassac for critical comments on the manuscript. We acknowledge funding by the Deutsche Forschungsgemeinschaft (grants BL462/15 to F.R.B. and HA7550/2 to D.H.).

## Additional information

### Funding

Deutsche Forschungsgemeinschaft BL462/15 Frank Blattner

Deutsche Forschungsgemeinschaft HA7550/2 Dörte Harpke

### Author contributions

Designed study: F.R.B., D.H. Coordinated study: Z.N., D.H., F.R.B. Provided data or materials: H.K. Performed experiments: Z.N. Analyzed data: Z.N., D.H., F.R.B., A.G. The initial manuscript was written by F.R.B. All authors contributed to and approved the final version of the manuscript.

### Data Accessibility

All sequence data are available through DDBJ/ENA/GenBank for the nuclear single-copy genes and the chloroplast locus (LS975036–LS975118), the annotated chloroplast genomes (MH542231–MH542233), and the GBS data (ERR2740826–ERR2740842, ERR2740845– ERR2741003).

## Additional files

### Supplementary files

**Figure supplement 1.** Parsimony consensus tree based on GBS sequence data.

**Figure supplement 2.** Maximum-likelihood phylogenetic tree based on GBS sequence data.

**Figure supplement 3.** Panel of Bayesian assignment analyses from K = 2 to K = 9.

**Figure supplement 4.** Chloroplast genome map for *C. cartwrightianus*.

**Figure supplement 5.** Panel of five single-copy gene trees calculated with the maximum-likelihood algorithm to infer the influence of allele diversity in *C. cartwrightianus* on the phylogenetic position of *C. sativus* and other included *Crocus* species.

**Figure supplement 6.** Photographs of *C. sativus* and three individuals of Attic *C. cartwrightianus*.

**Table supplement 1.** Studied plant materials.

**Table supplement 2.** Analyzed nuclear and chloroplast loci.

## Supplementary files

**Figure S1.**
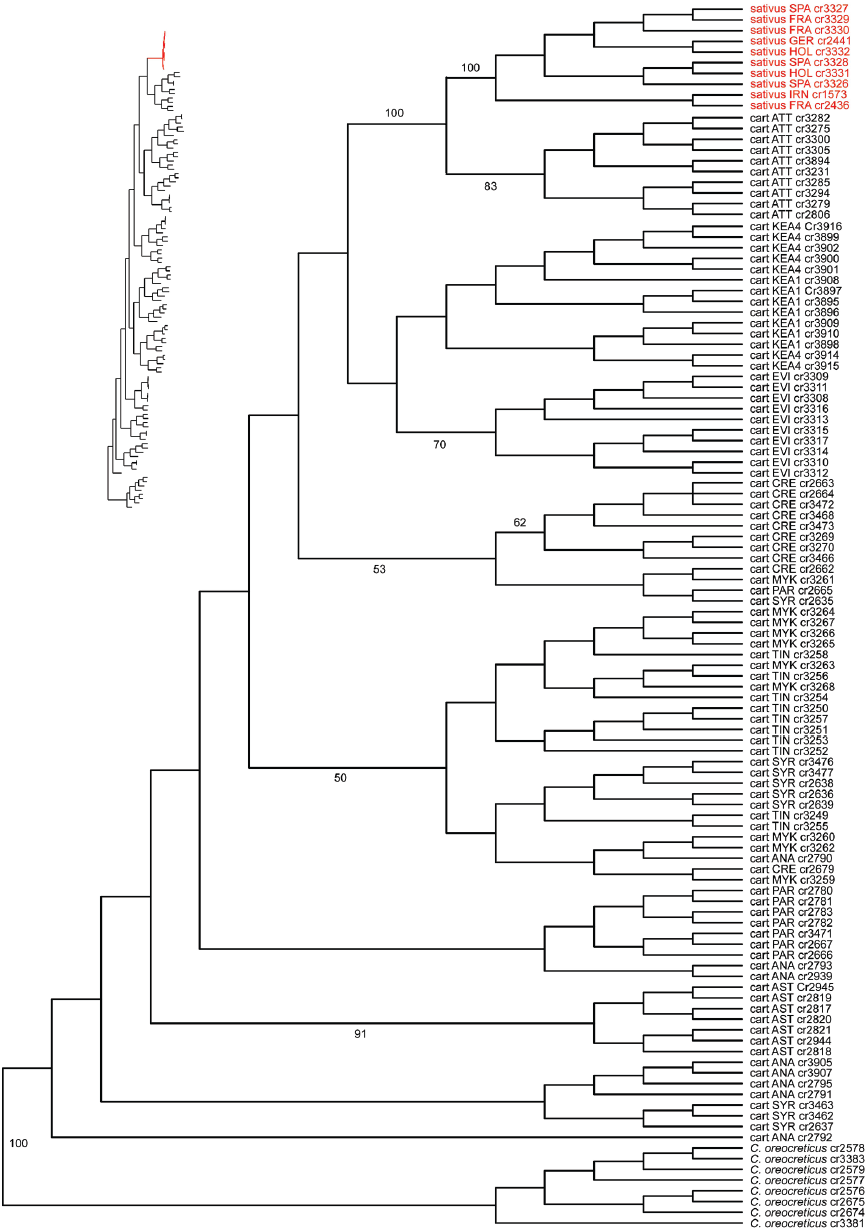
Maximum parsimony consensus tree of two most parsimonious trees obtained through the analysis of the GBS dataset. The inset on the left side provides branch lengths for one of the two trees. Numbers along branches depict bootstrap values ≥50% for major clades.

**Figure S2.**
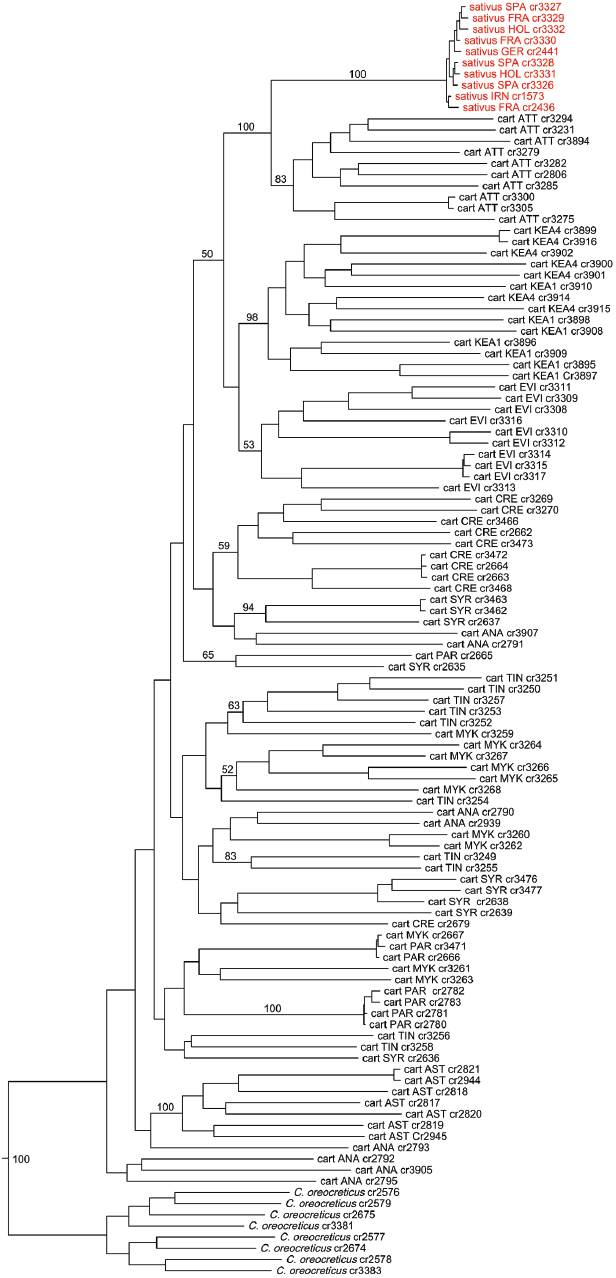
Maximum-likelihood tree obtained through the analysis of the GBS dataset with RAXML. Bootstrap values ≥50% are given for geographically defined clades.

**Figure S3.**
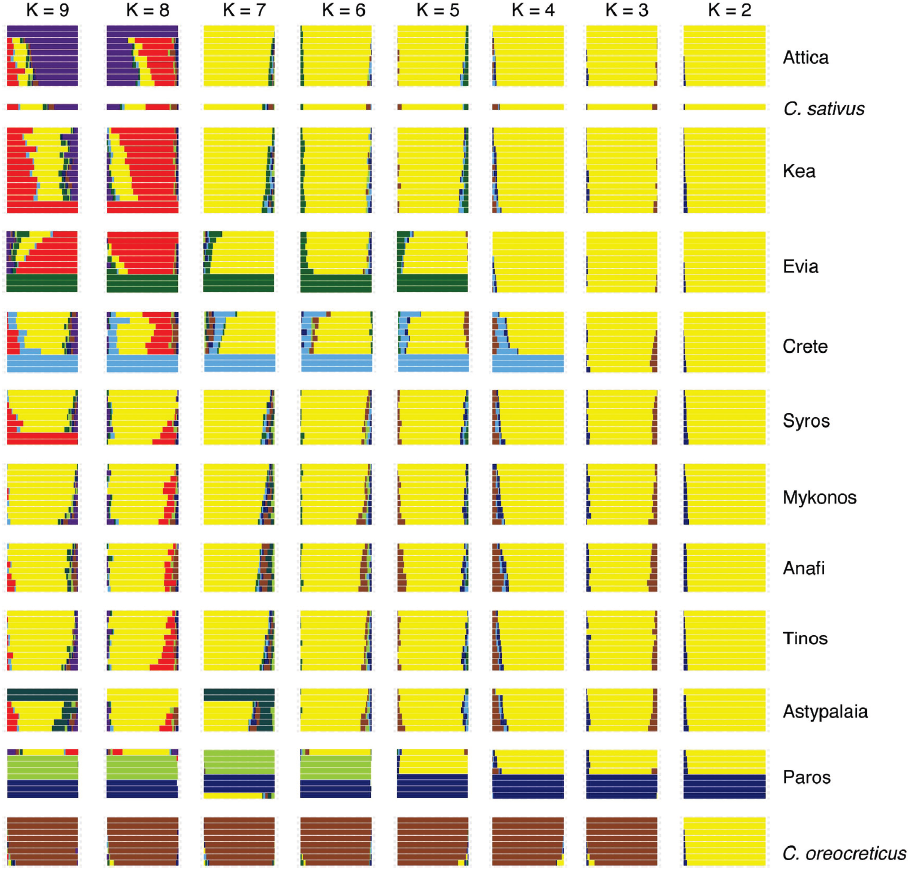
Bayesian assignment analysis plot for the GBS data with K = 2 to K = 9. *Crocus sativus* is assigned to the population from Attica although in some runs it shows also allele patterns similar to individuals from the island of Kea, however, with different proportions.

**Figure S4.**
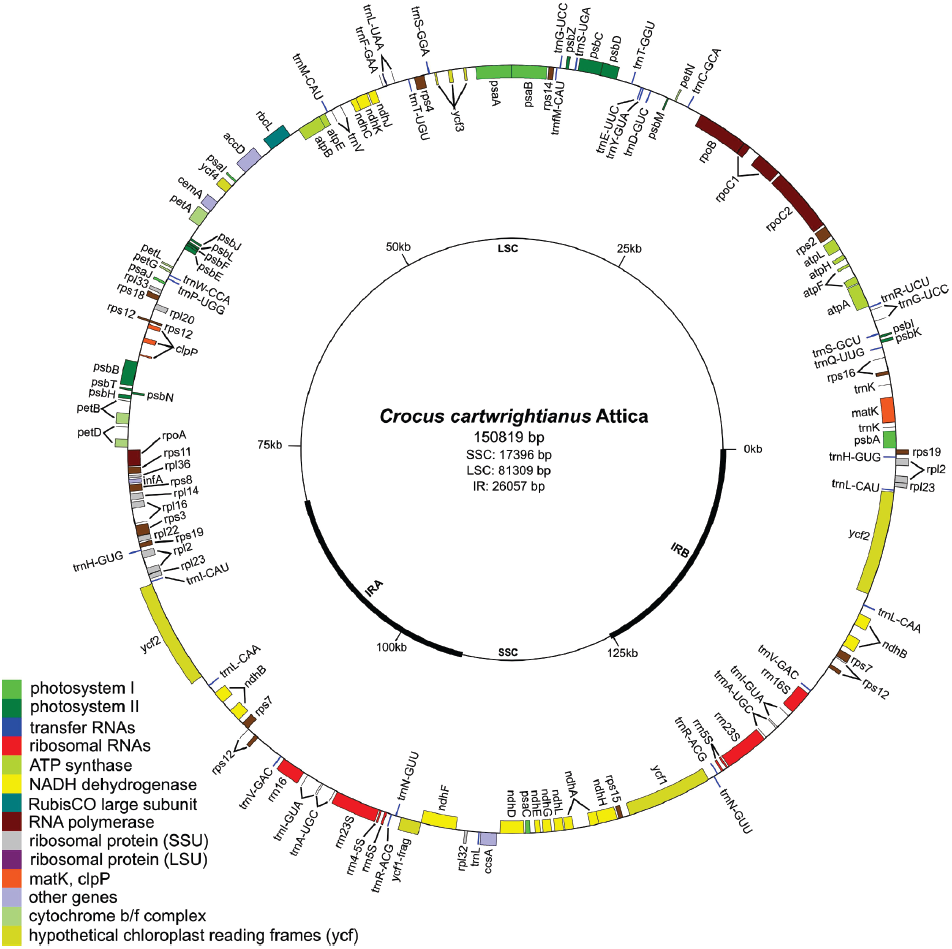
Annotated chloroplast genome of *C. cartwrightianus.* Two other assembled chloroplast genomes (*C. cartwrightianus* from Crete and *C. sativus*) are identical regarding gene order.

**Figure S5.**
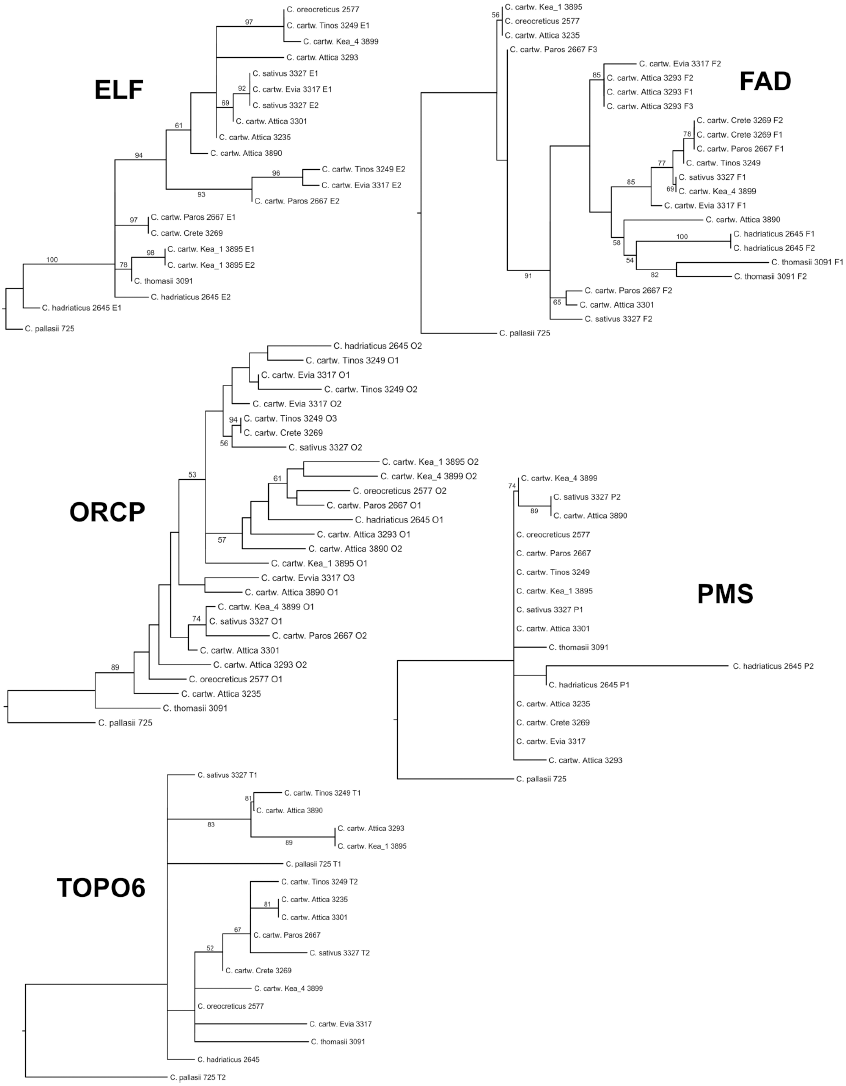
Maximum-liklihood analyses of five single-copy loci in close relatives of *Crocus sativus*. We analyzed multiple individuals of *C. cartwrightianus* to see how allelic diversity of the five genes influences the outcome of phylogenetic analyses regarding the position of *C. sativus* in relation to *C. cartwrightianus* and other included species. *Crocus pallasii* was specified as outgroup in all analyses. The presence of multiple copies of the genes was detected via cloning and sequencing of PCR amplicons. Different copies (paralogs or homeologs) present in single individuals are indicated by the first letter of the gene name followed by numbers 1-3. Numbers along branches depict bootstrap support values derived from 500 bootstrap re-samples. For description of the loci see Table S2.

**Figure S6.**
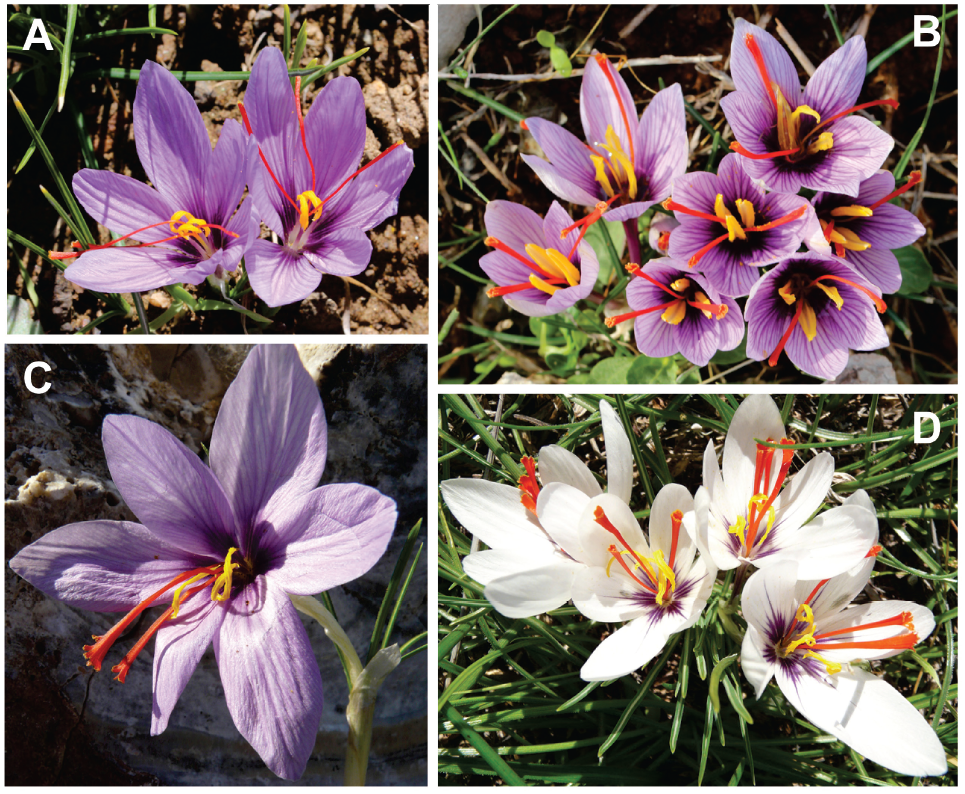
Comparison of *C. sativus* (A) and different individuals of Attic *C. cartwrightianus* (B-D).

**Table S1.**
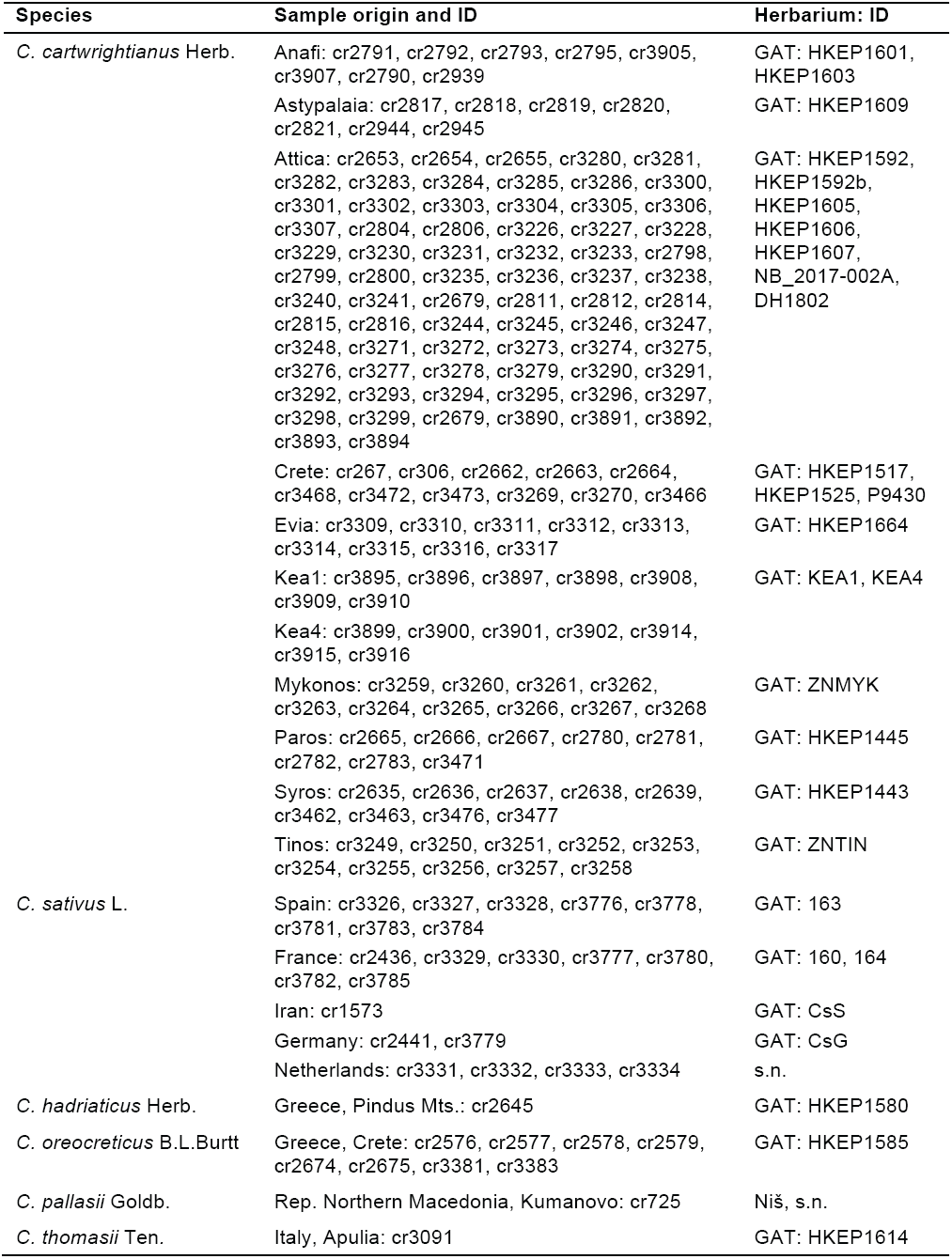
Plant materials used in the study.

**Table S2.**
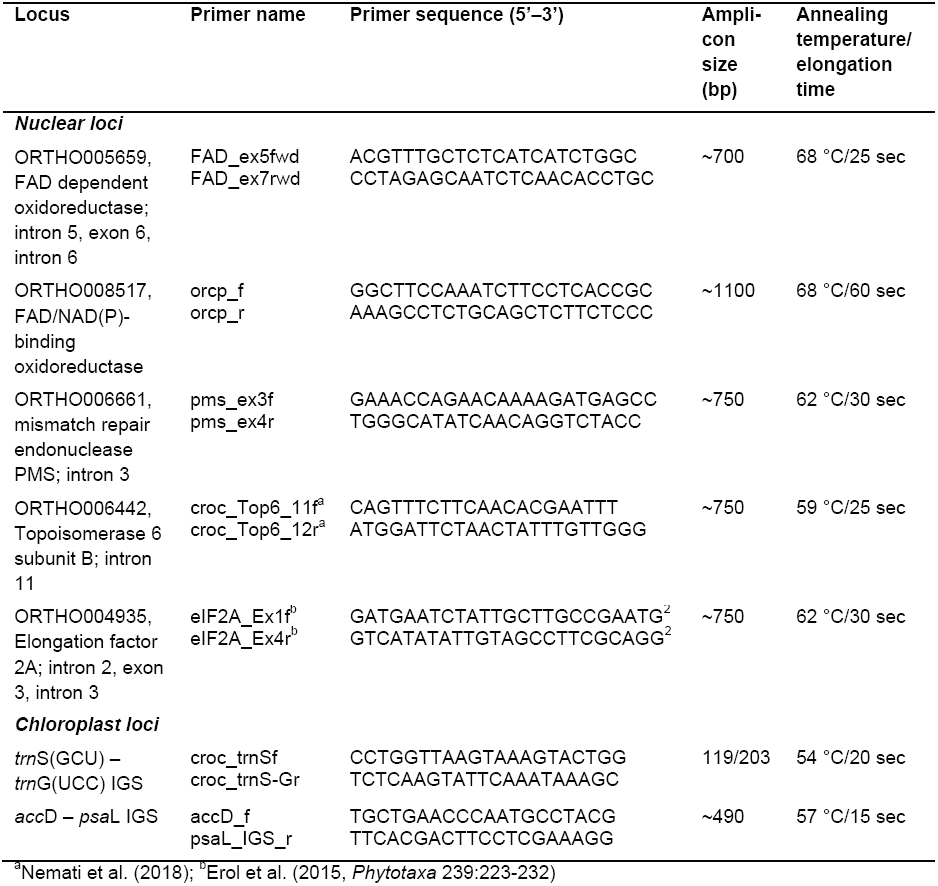
Analyzed genome regions.

